# Body size and composition and site-specific cancers in UK Biobank: a Mendelian randomisation study

**DOI:** 10.1101/2020.02.28.970459

**Authors:** Mathew Vithayathil, Paul Carter, Siddhartha Kar, Amy M. Mason, Stephen Burgess, Susanna C. Larsson

**Affiliations:** MRC Cancer Unit, University of Cambridge, Cambridge, UK; Department of Public Health and Primary Care, University of Cambridge, Cambridge, UK; MRC Integrative Epidemiology Unit, Bristol Medical School, University of Bristol, Bristol, UK; MRC Biostatistics Unit, University of Cambridge, Cambridge, UK; Unit of Cardiovascular and Nutritional Epidemiology, Institute of Environmental Medicine, Karolinska Institutet, Stockholm, Sweden; Department of Surgical Sciences, Uppsala University, Uppsala, Sweden

**Keywords:** cancer, body mass index, height, Mendelian randomization study, genomic, risk

## Abstract

**Objectives:** To investigate the casual role of body mass index, body fat composition and height in cancer.

**Design:** Two stage mendelian randomisation study

**Setting:** Previous genome wide association studies and the UK Biobank

**Participants:** Genetic instrumental variables for body mass index (BMI), fat mass index (FMI), fat free mass index (FFMI) and height from previous genome wide association studies and UK Biobank. Cancer outcomes from 367 586 participants of European descent from the UK Biobank.

**Main outcome measures:** Overall cancer risk and 22 site-specific cancers risk for genetic instrumental variables for BMI, FMI, FFMI and height.

**Results:** Genetically predicted BMI (per 1 kg/m^2^) was not associated with overall cancer risk (OR 0.99; 95% confidence interval (CI) 0-98-1.00, *p*=0.105). Elevated BMI was associated with increased risk of stomach cancer (OR 1.15, 95% (CI) 1.05-1.26; *p*=0.003) and melanoma (OR 0.96, 95% CI 0.92-1.00; *p*=0.044). For sex-specific cancers, BMI was positively associated with uterine cancer (OR 1.08, 95% CI 1.01-1.14; *p*=0.015) but inversely associated with breast (OR 0.95, 95% CI 0.92-0.98; *p*=0.001), prostate (OR 0.95, 95% CI 0.92-0.99; *p*=0.007) and testicular cancer (OR 0.89, 95% CI 0.81-0.98; *p*=0.017). Elevated FMI (per 1 kg/m^2^) was associated with gastrointestinal cancer (stomach cancer OR 4.23, 95% CI 1.18-15.13, *p*=0.027; colorectal cancer OR 1.94, 95% CI 1.23-3.07; *p*=0.004). Increased height (per 1 standard deviation, approximately 6.5cm) was associated with increased risk of overall cancer (OR 1.06; 95% 1.04-1.09; *p* = 2.97×10^-8^) and most site-specific cancers with the strongest estimates for kidney, non-Hodgkin lymphoma, colorectal, lung, melanoma and breast cancer.

**Conclusions:** There is little evidence for BMI as a casual risk factor for cancer. BMI may have a causal role for sex-specific cancers, although with inconsistent directions of effect, and FMI for gastrointestinal malignancies. Elevated height is a risk factor for overall cancer and multiple site cancers.

## BACKGROUND

Obesity is a global epidemic[1], with 20% of the world’s population predicted to be obese by 2025[2]. Multiple studies have demonstrated that obesity is associated with increased risk of cardiovascular[3][4], liver[5], and musculoskeletal disease[6]. The relationship between obesity and cancer risk has been subject to extensive investigation. Body mass index (BMI) is the most common measured marker for obesity, and correlates with fat mass[7]. Observational studies show raised BMI is associated with increased risk[8–11], no risk[8,11] and even reduced risk[8,12] of cancer. Adult height has been shown to be positively associated with multiple cancer types[13–15] and may also be a risk factor for carcinogenesis. However, traditional epidemiological studies are subject to confounding factors and reverse causation, and the true relationship between obesity and cancer remains unclear.

Mendelian randomisation (MR) uses genetic variants as instrumental variables for an exposure to assess the causal effect of the exposure on a disease outcome. As a result of Mendel’s laws of segregation and independent assortment, estimates from MR are less susceptible to bias due to confounding factors and reverse causality than those from conventional observational epidemiology. Alternative indices to BMI, such as fat mass index (FMI) and fat-free mass index (FFMI) can provide a more accurate measure of body fat composition[16,17][18], and further determine the specific contribution of adipose and non-adipose tissue to cancer risk.

In this study, we used MR to determine the causal role of genetically predicted BMI, FMI, FFMI and height on the risk of developing 22 cancers in 367 586 individuals from the UK Biobank database.

## METHODS

### Study population

Data for the genetic associations with site-specific cancer risk were obtained from UK Biobank. UK Biobank comprises of demographic, clinical, biochemical and genetic data from around 500 000 adults (aged 37-73 years old) recruited between 2006 and 2010 and followed up until 31st March 2017[19]. Only unrelated individuals of European descent (defined by self-report and genetics) were included in our analysis in order to reduce population stratification bias. After performing quality control filters as described previously[20], 367 586 individuals were included in analyses. We defined cancer outcomes in UK Biobank for the 22 most common site-specific cancers in the UK using ICD-9 and ICD-10 coding (Supplementary Table S1). Cancer outcomes were obtained from electronic health records, hospital episodes statistics (HES) data, the national cancer registry, death certification data and self-reporting validated by nurse interview. Genetic association estimates were obtained for each cancer outcome by logistic regression adjusting for age, sex and ten genetic principal components. Associations for sex-specific cancers (breast, cervical and uterine for women, testicular and prostate for men) were estimated in participants of the relevant sex only (198,838 women and 168,748 men).

### Genetic instruments

The genetic instrument for BMI in this study comprised 96 single-nucleotide polymorphisms (SNPs) (Supplementary Table S2) previously shown to be associated with BMI at genome-wide significance (*p* <5×10^-8^) in a genome-wide association study (GWAS) of 339 224 individuals of mainly (95%) European descent[21]. That GWAS identified 97 independent SNPs but one SNP was unavailable in UK Biobank. For FMI and FFMI (measured using bioelectrical impedance), the 98 SNPs associated with body composition among 362 499 UK Biobank participants[22] were considered. Out of those SNPs, 82 were in Hardy-Weinberg equilibrium (p>0.01) and had an imputation quality score above 0.8 and were included in these analyses (Supplementary Table S3). We computed FMI and FFMI by dividing fat mass or fat-free mass by the square of height. A GWAS of 253 288 European-descent individuals identified 697 genome-wide significant SNPs (*p* <5×10^-8^) for adult height[23] of which 690 were available in UK Biobank and used as instrumental variables (Supplementary Table S2).

### Statistical analysis

Associations of genetically-predicted BMI, FMI, FFMI and height with site-specific cancers were obtained using the random-effects inverse-variance weighted approach[24]. Analyses were also performed for BMI and height with overall cancer risk. We performed sensitivity analysis using the weighted median[24] and MR-Egger[25] methods. The analyses of FMI and FFMI were based on the multivariable inverse-variance weighted method with both exposures included in the same model. The odds ratios (OR) are expressed as per 1 kg/m^2^ increase in genetically predicted BMI, FMI, and FFMI and per 1 standard deviation (approximately 6.5cm) increase in height. As the number of cases and thus statistical power differed between analyses, we did not set a threshold for statistical significance. Statistical analyses were performed in Stata/SE 14.2 and R 3.6.0 software.

### Patient and Public Involvement

Patients and public users were not involved in the study design or analysis.

## RESULTS

367 586 participants were included in the study. The mean age was 57.2 years, with 46% males. 10.3% of participants were current smokers. In UK Biobank, the 96 SNPs explained 1.6% of the variance in BMI, whereas the 82 SNPs for body composition explained 0.8% of the variance in fat mass index and 0.7% of the variance in fat-free mass index. The 690 SNPs for height explained around 16% of the variance in height.

### BMI and Cancer Risk

A total of 75 037 cancer events were found in the UK Biobank. Genetically elevated BMI was not associated with overall cancer (OR 0.99; 95% confidence interval (CI) 0-98-1.00, *p*=0.105) but it was associated with an increased risk of stomach cancer (OR 1.15, 95% confidence interval (CI) 1.05-1.26; *p*=0.003) (Figure 1). For sex-specific cancers, elevated BMI was positively associated with uterine cancer (OR 1.08, 95% CI 1.01-1.14; *p*=0.015) but inversely associated with breast (OR 0.95, 95% CI 0.92-0.98; *p*=0.001), prostate (OR 0.95, 95% CI 0.89-1.02; *p*=0.007), and testicular (OR 0.89, 95% CI 0.81-0.98; *p*=0.017) cancers. An inverse association between BMI and melanoma was observed (OR 0.96, 95% CI 0.92-1.00; *p*=0.044). Results were similar but less precise in sensitivity analyses (Supplementary Table S4).

**Figure 1.**
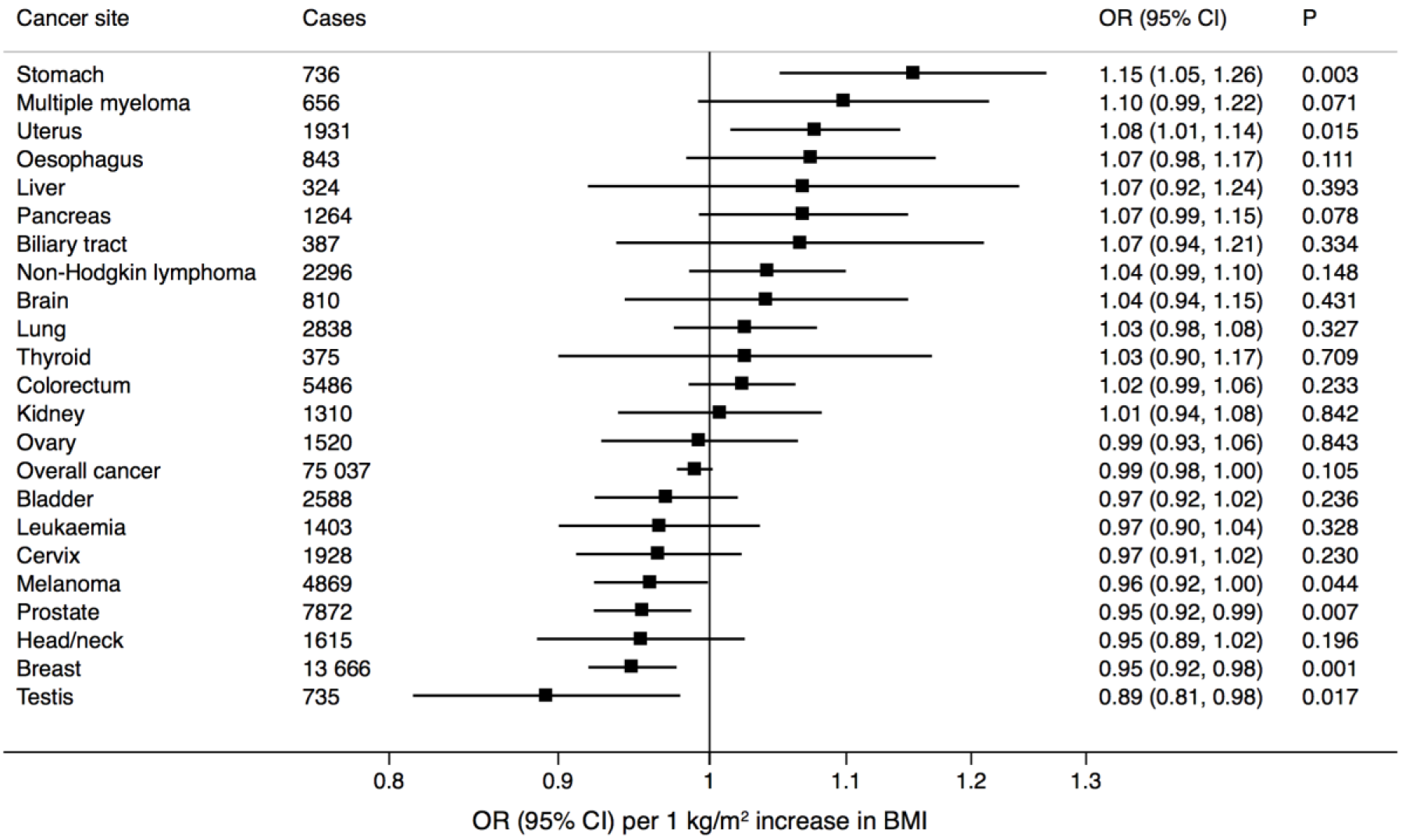
Associations of genetic predisposition to higher body mass index (BMI) with site-specific cancers. Odds ratios are per 1 kg/m^2^ increase in BMI. Results are obtained from the random-effects inverse-variance weighted method. CI, confidence interval; OR, odds ratio.

### FMI and FFMI and Cancer Risk

No associations were observed between overall cancer risk and elevated FMI (OR 1.09, 95% CI 0.93-1.28; *p*=0.297) and FFMI (OR 0.90, 95% CI 0.77-1.05; *p*=0.197). Similar to BMI, genetically raised FMI was associated with risk of stomach cancer (OR 4.23, 95% CI 1.18-15.13; *p*=0.027) (Figure 2). Additionally, a positive association was observed with colorectal cancer (OR 1.94, 95% CI 1.23-3.07; *p*=0.004). Suggestive positive associations were observed with liver (OR 4.28, 95% CI 0.67-27.20; *p*=0.124) and lung cancer (OR 1.79, 95% CI 0.95-3.36; *p*=0.072). Genetically elevated FFMI had an inverse association with head and neck cancer (OR 0.48, 95% CI 0.23-0.98; *p*=0.044), and a suggestive inverse association with colorectal cancer (OR 0.63, 95% CI 0.42-1.00; *p*=0.050) (Figure 3).

**Figure 2.**
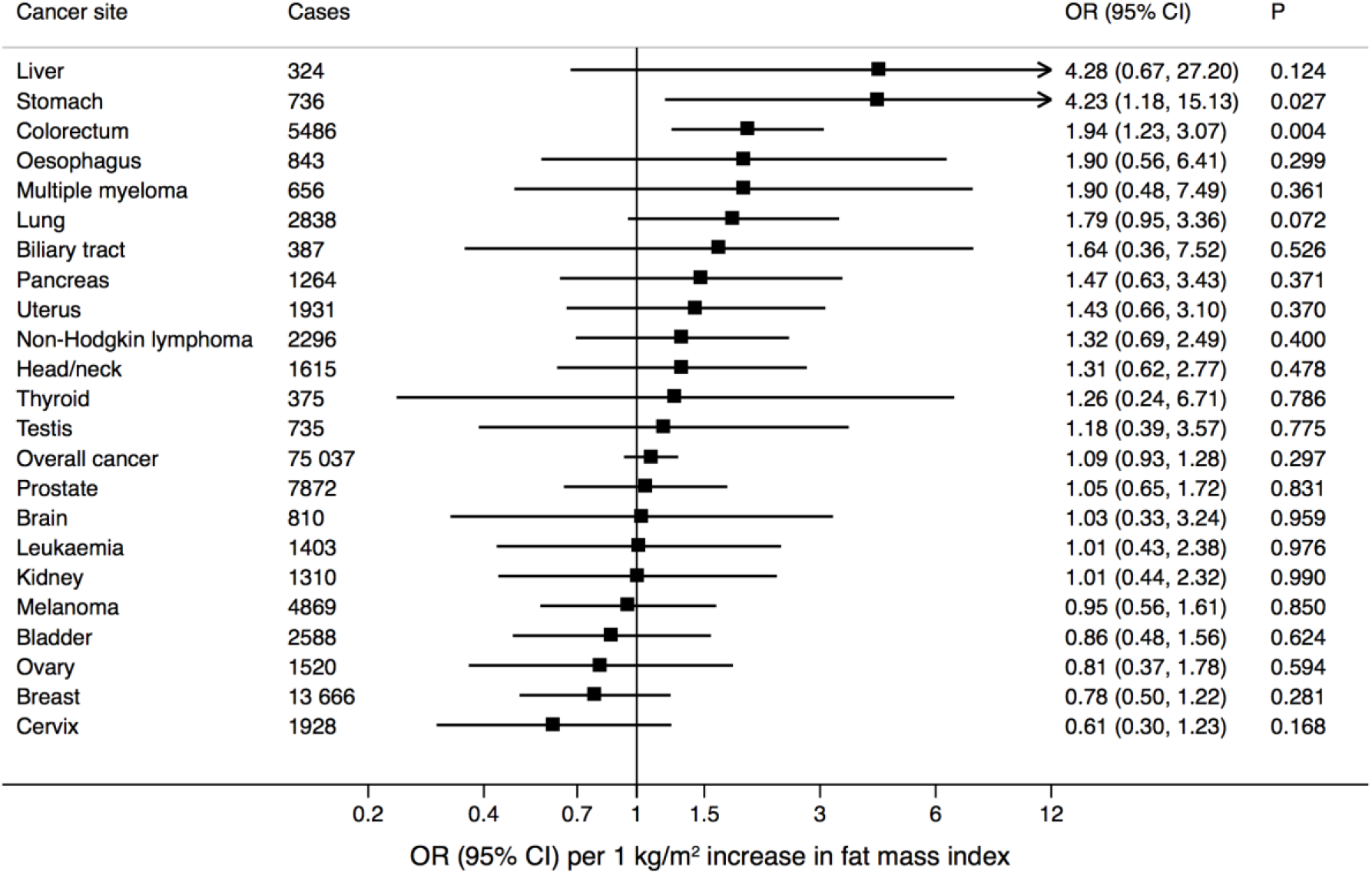
Associations of genetic predisposition to higher fat mass index (FMI) with site-specific cancers. Odds ratios are per one 1 kg/m^2^ increase in FMI. Results are obtained from the random-effects inverse-variance weighted method. CI, confidence interval; OR, odds ratio.

**Figure 3.**
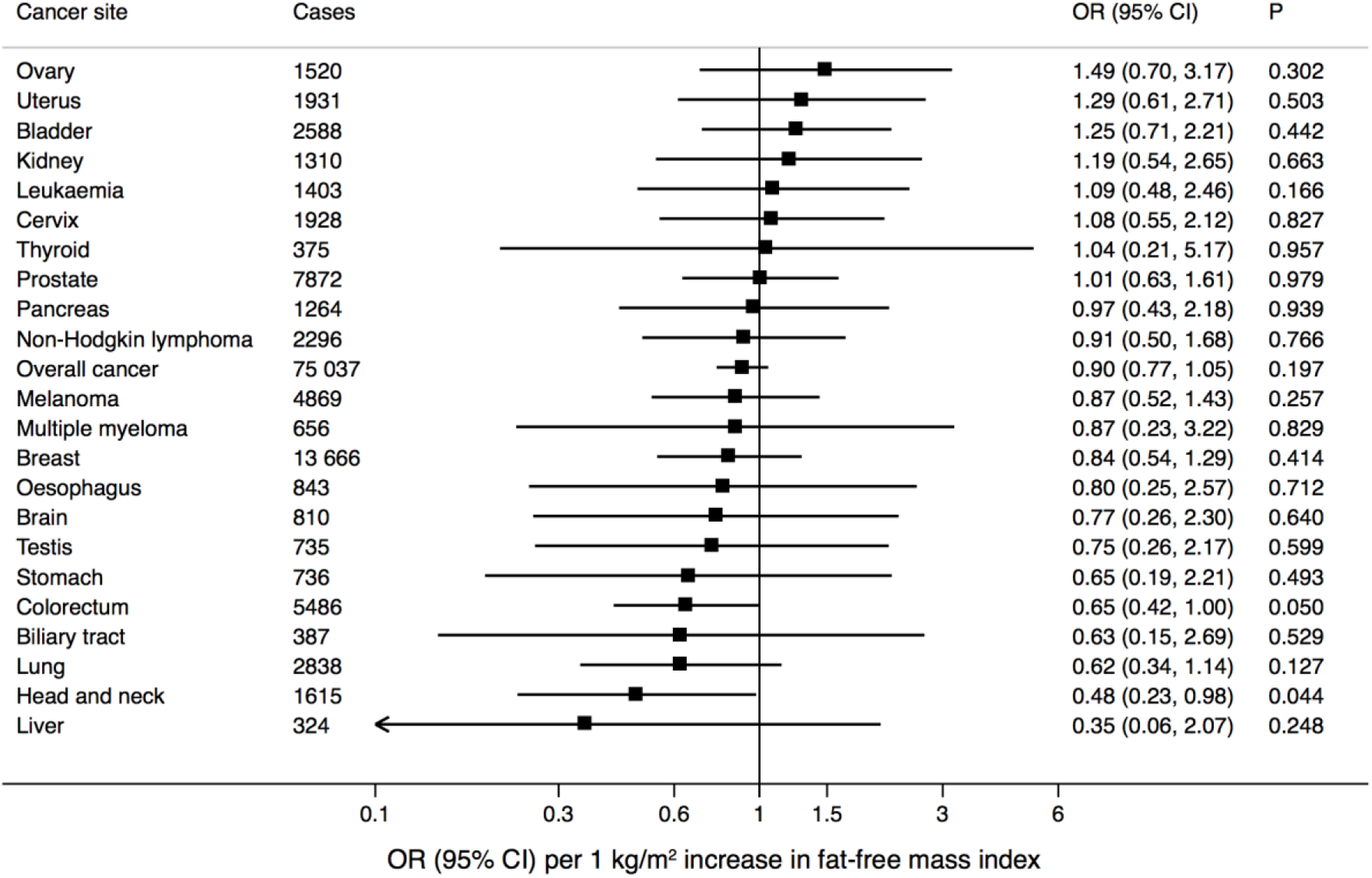
Associations of genetic predisposition to higher fat-free mass index (FFMI) with site-specific cancers. Odds ratios are per 1 kg/m^2^ increase in FFMI. Results are obtained from the random-effects inverse-variance weighted method. CI, confidence interval; OR, odds ratio.

### Height and Cancer Risk

Elevated height was associated with overall cancer (OR 1.06; 95% 1.04-1.09; *p* = 2.97×10^-8^) and multiple site-specific cancers, including kidney (OR 1.14, 95% CI 1.01-1.29; *p*=0.034), non-Hodgkin lymphoma (OR 1.12, 95% CI 1.03-1.23; *p*=0.012), colorectal (OR 1.12, 95% CI 1.05-1.19; *p*=0.001), lung (OR 1.11, 95% CI 1.02-1.20; *p*=0.013), melanoma (OR 1.09, 95% CI 1.02-1.17; *p*=0.014) and breast cancer (OR 1.06, 95% CI 1.01-1.11; *p*=0.012) (Figure 4). There were also suggestive associations of height with cancers of the liver (OR 1.26, 95% CI 0.98-1.62; *p*=0.067), biliary tract (OR 1.24; 95% CI 0.99-1.54; *p*=0.062) and thyroid (OR 1.23, 95% CI 0.99-1.54; *p*=0.064). The associations remained in sensitivity analyses (Supplementary Table S5).

**Figure 4.**
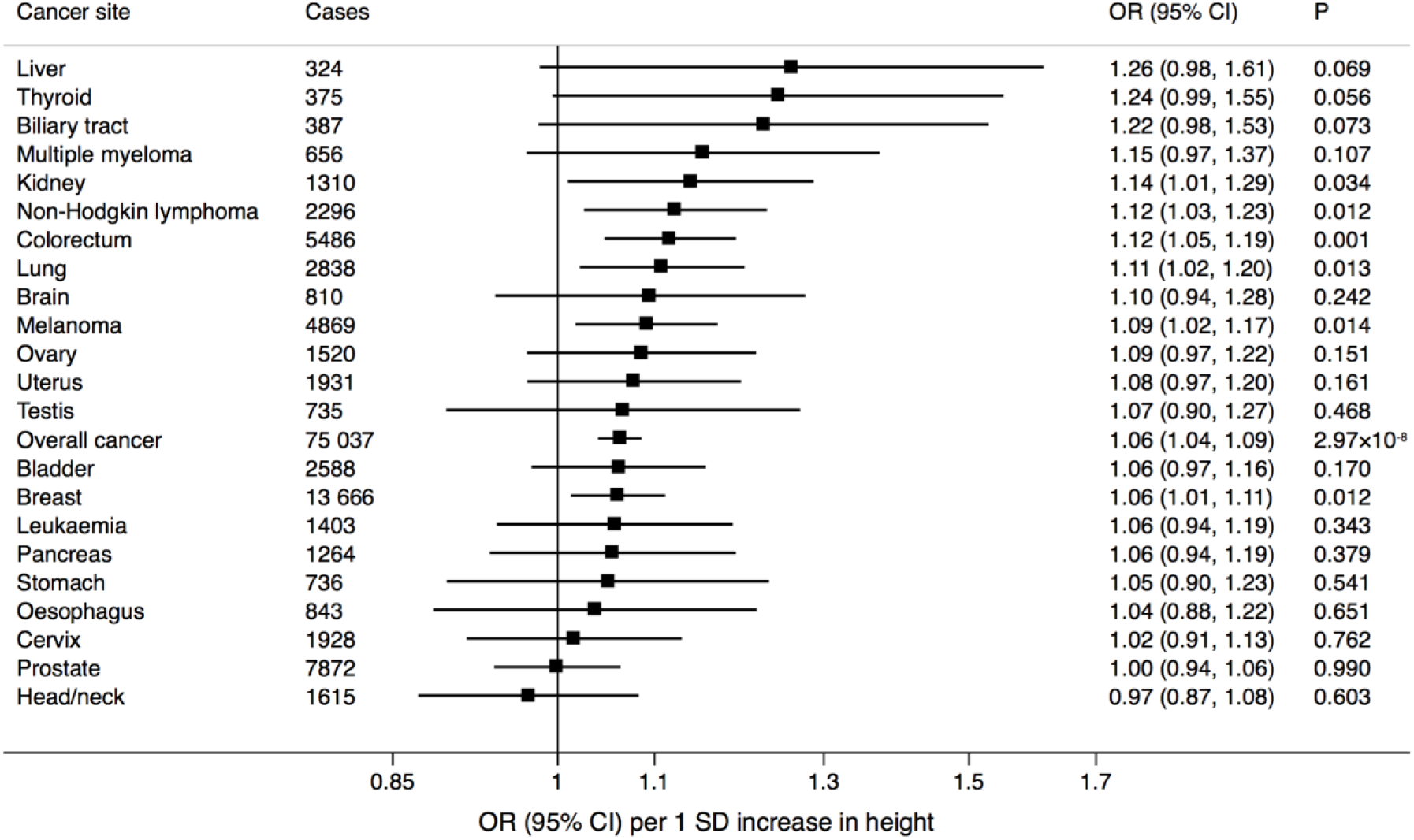
Associations of genetic predisposition to increased height with site-specific cancers. Odds ratios are per 1 standard deviation (6.5cm) increase in height. Results are obtained from the random-effects inverse-variance weighted method. CI, confidence interval; OR, odds ratio.

## DISCUSSION

In the present MR study of 367 586 individuals, we found that genetically predicted BMI was not associated with overall cancer risk but positively associated with stomach and uterine cancer, and inversely associated with breast, prostate, testicular cancer and melanoma. FMI was positively associated with stomach and colorectal cancer. FFMI showed inverse associations with head and neck cancer and colorectal cancer. Height was positively associated with overall and several site-specific cancers.

The association between BMI and adiposity with risk of certain gastrointestinal cancers which we report replicates and extends previous findings. Colorectal cancer has been positively associated with BMI in a meta-analysis of 23 observational studies[9] as well as with waist-to-hip ratio (WHR)[26] and BMI[27,28] in previous MR analyses. Similarly, high BMI has been associated with gastric cancer in a meta-analysis of observational studies[29] and in an MR study of 2631 Chinese adults[30]. We support these findings and suggest a causal role of adiposity in driving certain cancers with the observations that elevated FMI was associated with stomach and colorectal cancer, with the latter inversely associated with FFMI. Mechanisms by which adipose tissue could drive gastrointestinal carcinogenesis have previously been studied. Increased adipose tissue is associated with insulin resistance, and hyperinsulinaemia[31], with raised circulating insulin enhancing colorectal epithelial cell proliferation in rat models[32]. Additionally, ghrelin is a gut hormone produced in the stomach, with reduced levels seen in obesity[33]. Ghrelin reduces pro-inflammatory cytokines and inflammatory stress[34] and reduced levels are associated with increased risk of colorectal[35] and stomach[36] cancers. Adiposity is also well established in causing non-alcoholic fatty liver disease and has been implicated in its progression to hepatocellular carcinoma[37]. In line with this, we observed a low precision estimate for an increased risk of liver cancer with raised FMI. Our results therefore suggest causal roles between adiposity and certain gastrointestinal cancers and further research should assess the impact of weight loss interventions on this risk.

In this MR study, raised BMI was associated with sex-specific cancers with increased risk of uterine cancer, and decreased risk of breast, prostate and testicular cancer. BMI and breast cancer has been extensively studied in previous MR studies[28,38–40]. A large MR study based on data from the BCAC and DRIVE consortia of 46,325 cases of breast cancer found that 84 BMI-related SNPs were associated with reduced breast cancer risk in both pre- and post-menopausal women[39], consistent with the findings of our study. Furthermore, our study corroborates the findings of a previous MR study of 6,609 cases of uterine cancer, showing BMI was positively associated with incidence[41]. In males, our findings are in line with a previous MR of 22 European cohorts which showed a non-significant inverse association between BMI and prostate cancer[42]. Previous epidemiological studies have also shown an inverse association between BMI and testicular cancer[43], although we are the first to suggest a causal role through MR analysis. The association of BMI and these reproductive cancers is likely to be at least in part hormonally-mediated. In pre-menopausal women, increased BMI is associated with anovulation, reducing lifetime exposure of circulating oestrogen and progesterone[44], thus lowering breast cancer risk[45] and increasing uterine cancer risk[46]. Higher maternal BMI is also associated with reduced oestrogen exposure in utero[47], which is protective against testicular cancer[48] and in males, elevated BMI reduces serum testosterone[49], reducing prostate cancer risk. Our findings of causal roles of BMI in regulating risk of certain reproductive and hormone-responsive cancers are therefore biologically plausible although the mechanistic links need to be assessed further.

We observed a positive association between height and overall cancer risk, which was consistent across a wide range of site specific cancers: kidney, lymphoma, colorectal, lung, melanoma, breast. Additionally, there was suggestive evidence of positive associations between height and liver, biliary tract and thyroid cancers. Our findings are consistent with previous MR studies showing a positive association of height with colorectal[50,51], lung[50,52] and breast[53,54] cancer. Increased height is associated with elevated insulin-like growth factor 1 (IGF1),[55] which is a growth factor that drives cellular proliferation and survival and has thus been implicated in carcinogenesis of IGF responsive tissues. Notably therefore, and increased expression of IGF1 and its cellular receptors are present in cancer[56,57]. Dysregulation of IGF1 signalling axis in taller individuals may therefore be a driver in a wide range of cancers. Therapeutic options exist with receptor tyrosine kinase inhibitors and monoclonal inhibitors, although clinical trials using such agents in treatment of cancers have thus far had disappointing results[57]. Further studies should strengthen the link between IGF-1 in cancer progression in taller individuals and the impact of such therapies in taller patients with IGF responsive tumours.

The results from our MR analysis have clinical implications. Overall obesity is not associated with increased risk of overall cancer. This challenges recommendations from the International Agency for Cancer Research (IACR)[58] and CRUK[59] which present obesity as a principal cause of cancer. These conclusions are based on traditional observational studies which are subject to confounding and reverse causality, which are minimized in this MR design. Though a recent MR analysis showed obesity is a key risk factor for cardiovascular disease[3], our study shows obesity is not a consistent risk factor for cancer. This has future implications for public health measures for cancer prevention and screening. However, we do support a causal role of obesity in driving and protecting against certain cancers, which suggests differential effects of BMI and obesity in different malignancies which should be explored further.

Our study has several strengths. The MR design minimizes the influence of environmental confounding factors and reverse causality, allowing for causal relationships to be better elucidated. The UK Biobank is a large prospective cohort, allowing multiple cancer types to be studied from a single dataset and comparisons of estimates across cancers to be made. However, there are some limitations. The main limitation is that estimates for lower frequency cancer types are subject to low precision and therefore results should be interpreted based on the magnitude of the associations rather than on *p* values alone. Another shortcoming is that our findings may not be applicable to other ethnic groups as we confined the study population to European descent individuals to minimize bias from population stratification bias. The UK Biobank may be subject to selection bias, and as such cancers, particularly those of younger ages, may not be captured. Furthermore, our study does not provide understanding of the physiological pathways in which obesity and height may affect carcinogenesis.

## CONCLUSION

In conclusion, this comprehensive MR study does not support a causal role between obesity and risk of overall cancer. However, we provide evidence that increased BMI, in particular elevated fat mass, increases risk of gastrointestinal and uterine cancer, but protective for other sex-specific cancers. Increased height increased the risk of overall and site specific cancers. Our findings challenge the dogma of obesity causing cancer.

## Supporting information

the UK using ICD-9 and ICD-10 coding (Supplementary Table S1).

## ACKNOWLEDGMENTS

The authors thank all investigators from the UK Biobank, where data were conducted under application 29202. The views expressed are those of the authors and not necessarily those of the National Health Service, the National Institute for Health Research or the Department of Health and Social Care.

## ETHICAL APPROVAL

The UK Biobank received ethical approval from the research ethics committee (REC reference for UK Biobank 11/NW/0382). All participants provided written informed consent. All genetic data used is publically available and therefore no ethics approval was required.

## CONTRIBUTORS

MV, PC, SK, SB and SCL designed the investigation. SB, AM and SCL conducted data extraction and analysis. MV, PC, SK, AM, SB and SCL all contributed to interpretation of results and writing of the final manuscript.

